# Generating Electricity from Water Evaporation Through Microbial Biofilms

**DOI:** 10.1101/2021.12.14.472618

**Authors:** Xiaomeng Liu, Toshiyuki Ueki, Hongyan Gao, Trevor L. Woodard, Kelly P. Nevin, Tianda Fu, Shuai Fu, Lu Sun, Derek R. Lovley, Jun Yao

## Abstract

Sustainable strategies for energy production are required to reduce reliance on fossil fuels and to power electronics without generating toxic waste.^1-7^ Generating electricity from water evaporation through engineered materials is a promising approach,^8,9^ but power outputs have been low and the materials employed were not sustainably produced. Microorganisms can be mass produced with renewable feedstocks. Here, we demonstrate that it is possible to engineer microbial biofilms as a cohesive, flexible material for long-term continuous electricity production from evaporating water. The biofilm sheets were the functional component in devices that continuously produced power densities (∼1 μW/cm^2^) higher than that achieved with non-biological materials. Current production scaled directly with biofilm-sheet size and skin-patch devices harvested sufficient electricity from the moisture on skin to continuously power wearable devices. The results demonstrate that appropriately engineered biofilms can perform as robust functional materials without the need for further processing or maintaining cell viability. Biofilm-based hydroelectric current production was comparable to that achieved with similar sized biofilms catalyzing current production in microbial fuel cells,^10,11^ without the need for an organic feedstock or maintaining cell viability. The ubiquity of biofilms in nature suggests the possibility of additional sources of biomaterial for evaporation-based electricity generation and the possibility of harvesting electricity from diverse aqueous environments.

---

Water evaporating at water-solid interfaces can drive charge transport for electricity generation.^8,9^ In order to be effective, this streaming mechanism requires a material with a large surface area with associated mobile surface charges. Practicality will require low-cost materials and a minimum of processing. However, to date evaporation-based electricity generation has primarily focused on devices fabricated from thin films of assembled nanomaterials to improve surface area, and further functionalized to introduce hygroscopic surface groups.^8,9^ Furthermore, power densities and stability have been low. For example, an initial device based on functionalized carbon black achieved an energy density in the range of tens of nW/cm^2^ for days.^12^ A silicon-nanowire material increased power density to the µW/cm^2^ level,^13^ but the increased material cost and lower stability (<12 h) reduced practicality. The use of biomaterial such as wood exploits its innate porous structure and built-in hygroscopic groups for ease in fabrication,^14^ with the benefit of reducing production wastes associated with conventional inorganic materials. However, the energy density remains limited to nW/cm^2^ level, and the rigid and bulk form further limits scalable integration and wearable implementation.

*Geobacter sulfurreducens* strain CL-1 is a genetically modified strain that produces highly cohesive, electrically conductive biofilm sheets (< 100 µm thick) that could potentially serve as a sustainably produced, functional biopolymer.^15^ In order to determine whether the biofilm sheets were capable of evaporation-based current generation, the material was produced as previously described^15^ (Supplementary Fig. S1) yielding cohesive biofilm sheets ∼40 µm thick (Fig. S2). The sheets (Fig. 1a) were patterned for device fabrication with standard thin-film laser writing (Fig. 1b; Fig. S3). Transmission electron microscopy (TEM) confirmed cells within an extracellular polymeric matrix^16^ with nanofluidic channels (width, 100-500 nm) between the cells (Fig. 1c), as previously reported.^15^

**Figure 1.**
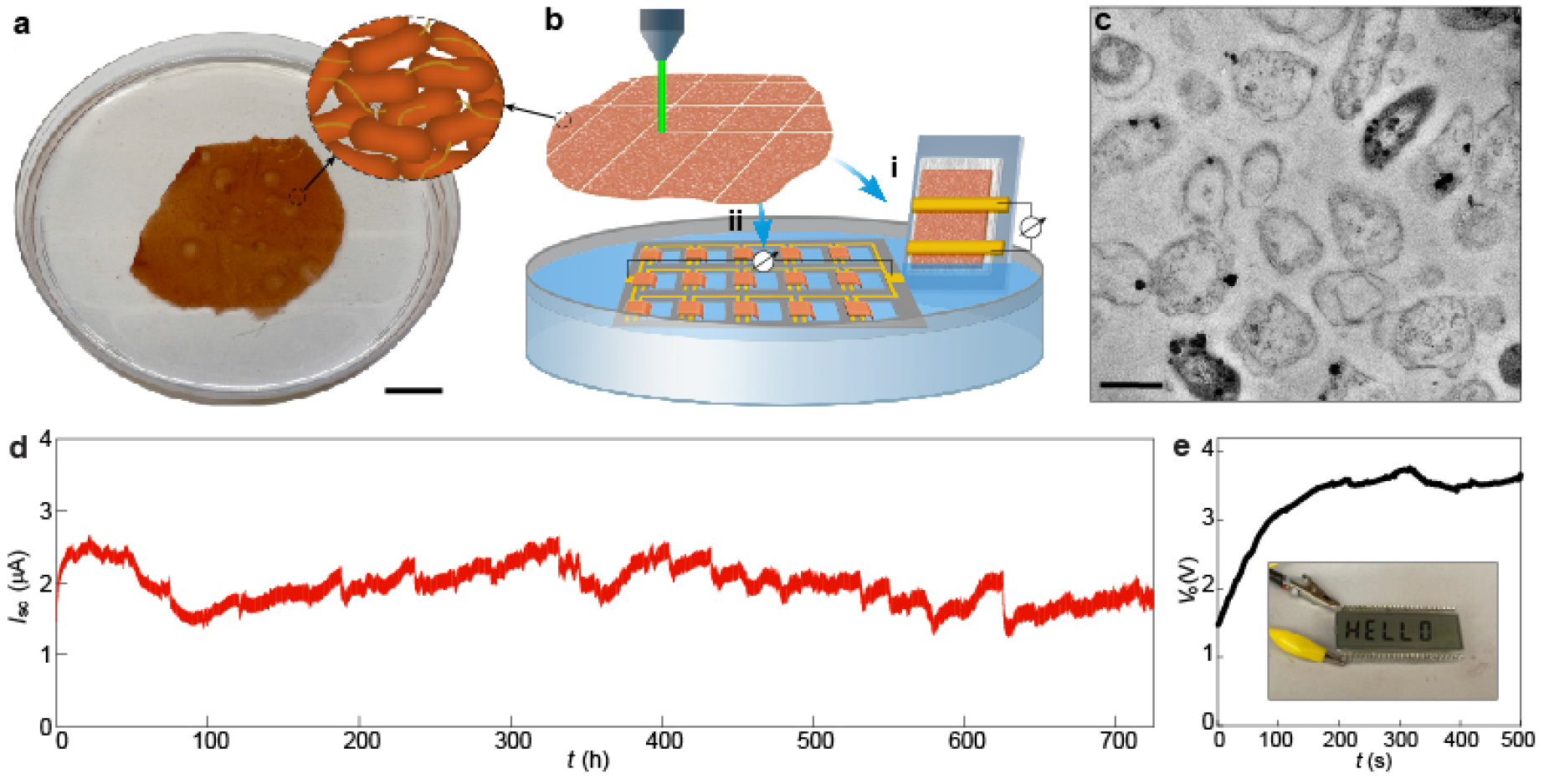
Electric outputs from *G. sulfurreducens* biofilms. **a**, A harvested biofilm sheet of *G. sulfurreducens* strain CL-1 (schematic inset), floating on water. Scale bars, 2 cm. **b**, Schematic of using laser-patterned biofilms to construct (i) single device and (ii) interconnected device array, with a portion of the biofilm at one electrode immersed in water. A tissue paper was used to support the biofilm in the single device, which may assist water evaporation but does not contribute to electric signals (supplementary Fig. S4). **c**, Cross-sectional TEM of a *G. sulfurreducens* strain CL-1 biofilm. Scale bar, 1 µm. **d**, A continuous recording of the short-circuit current (*I*_sc_) from a device for one month. The device had the structure shown in (b-i), with an electrode spacing 2 mm and lateral width 1 cm, yielding an estimated energy density *V*_o_·*I*_sc_/4 of ∼1 µW/cm^2^ in the active biofilm region. The test was performed in the ambient environment with a relative humidity (RH) fluctuating between 30-55%. **e**, Open-circuit voltage (*V*_o_) from an integrated device array (b-ii) floating on water surface, which was used to power up an LCD (inset). The actual device is shown in supplementary Fig. S8.

Electricity harvesting from water evaporation was initially evaluated in a device fabricated by placing two gold electrodes across a biofilm supported on a glass substrate (Fig. 1b-i; Fig. S4). Immersing one terminal in water yielded a spontaneous voltage output of ∼0.45 V, which was sustained for more than a month (Fig. S4). The electric outputs were the same when the inert gold electrodes were replaced with carbon electrodes (Fig. S5). Depleting the water source depleted the energy output (Fig. S6). Increasing the evaporation rate led to an increase in energy output (Fig. S7). These results are consistent with an evaporation-based streaming mechanism.^8,9^ The device maintained a steady current output (∼1.5 µA) for over 30 days (Fig. 1d). The estimated power density generated (∼1 μW/cm^2^) was more than an order of magnitude larger than that previously achieved with carbon and wood materials.^12,14^ Power output increased linearly when the biofilm-sheet devices were integrated in arrays (Fig. 1b-ii; Fig. S8), sufficient to power up electronics (Fig. 1e).

A vertical thin-film device was constructed to improve integration for scalable power production. The biofilm was sandwiched between a pair of mesh electrodes^17^ (Fig. 2a; Fig. S9) and the device was sealed with a pair of polydimethylsiloxane (PDMS) thin layers for structural stability. Placing the device on water surface resulted in a spontaneous voltage output ∼ 0.45 V (Fig. 2b, top), consistent with a vertical streaming potential derived from water evaporation across the film. The biofilm established the streaming potential over a short distance along both the vertical thickness and in the planar direction (Fig. S10), consistent with an isotropic transport expected from the 3D distributed porosity in the biofilm (Fig. 1c). This led to an enhanced electrical field, which was consistent with observed increase in power density. Connecting the two mesh electrodes yielded continuous current output (Fig. 2b, bottom).

**Figure 2.**
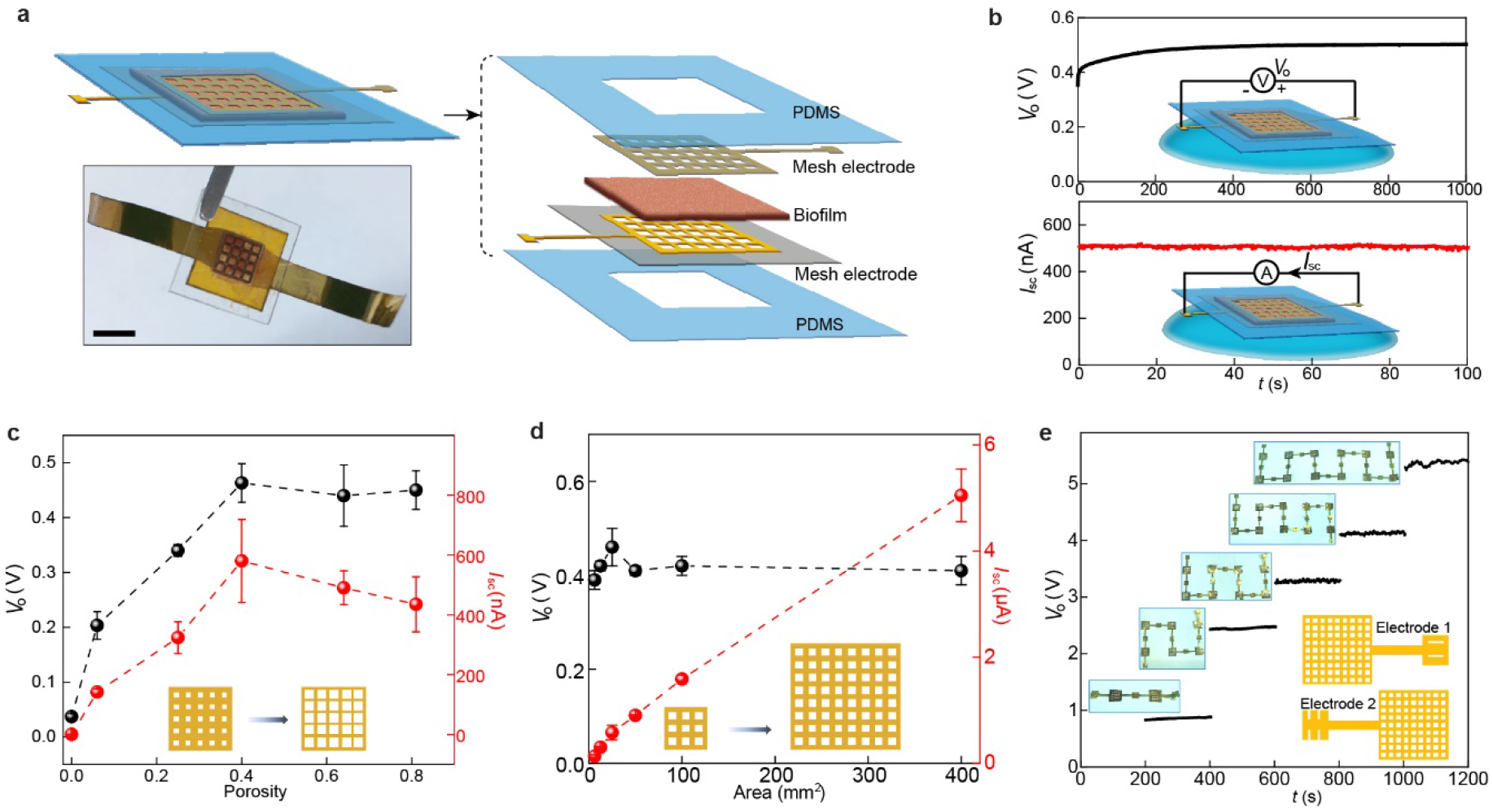
Integrated biofilm devices using mesh electrodes. Schematic and actual photo (bottom) of a biofilm device. Scale bar, 0.5 mm. **b**, Representative open-circuit voltage *V*_o_ (top) and short-circuit current *I*_sc_ (bottom) from a device placed on water surface (schematics). The device size was 5×5 mm^2^. **c**, *V*_o_ (black) and *I*_sc_ (red) from devices with respect to different porosities in the mesh electrodes. The device size was 25 mm^2^. The estimated optimal energy density (*V*_o_·*I*_sc_/4) was ∼0.7 µW/cm^2^ in the active biofilm region. The previous device structure (Fig. 1b-i) yielded higher density, probably because the biofilm-tissue paper interface improves water transport. **d**, *V*_o_ (black) and *I*_sc_ (red) from devices of different areas, with the porosity in the mesh electrodes kept 0.4. **e**, Measured *V*_o_ from devices connected in series (upper inset) using a ‘buckle’ design in electrodes (bottom schematic). All the tests were performed in the ambient environment (RH∼50%).

Increasing the porosity of the electrodes increased the voltage and current output up to a porosity of 0.4, with no additional benefit from greater porosity increases (Fig. 2c). The saturation trend in voltage is consistent with the concept that the limiting factor in the rate of water transport determining the streaming potential is eventually removed beyond a certain porosity. The current output gradually decreased at porosities above 0.4 (Fig. 2c, red), indicating that upon voltage saturation any further increase in porosity reduces the contact area for charge collection. There was a linear increase in current output with increase in device size, while the voltage remained constant (Fig. 2d). Devices were integrated for a tunable voltage output with a ‘buckle’ design that provided a rapid connect/disconnect (insets, Fig. 2e; Fig. S11). The output voltage linearly increased with an increase in the number of devices, with a voltage of ∼5 V readily obtained by connecting 12 devices in series (Fig. 2e). The biofilm-sheet arrays can be also integrated on the same flexible substrate to increase the output (Fig. S12).

High salt concentrations decrease the power output of some evaporation-based current generation devices^12^ because the decreased Debye length at high ionic strength reduces the double-layer overlap and hence the streaming efficiency.^18^ However, the biofilm-sheet devices maintained electric outputs in salt solutions with an ionic concentration (0.5 M) higher than that (∼0.15 M) in bodily water (Fig. 3a). This is attributed to the porous structure of the biofilm sheets (Fig. 1c) comprised of various organic groups,^16^ which can increase the effective Debye length,^19,20^ and hence, maintain the streaming efficiency at high ionic strength.

**Figure 3.**
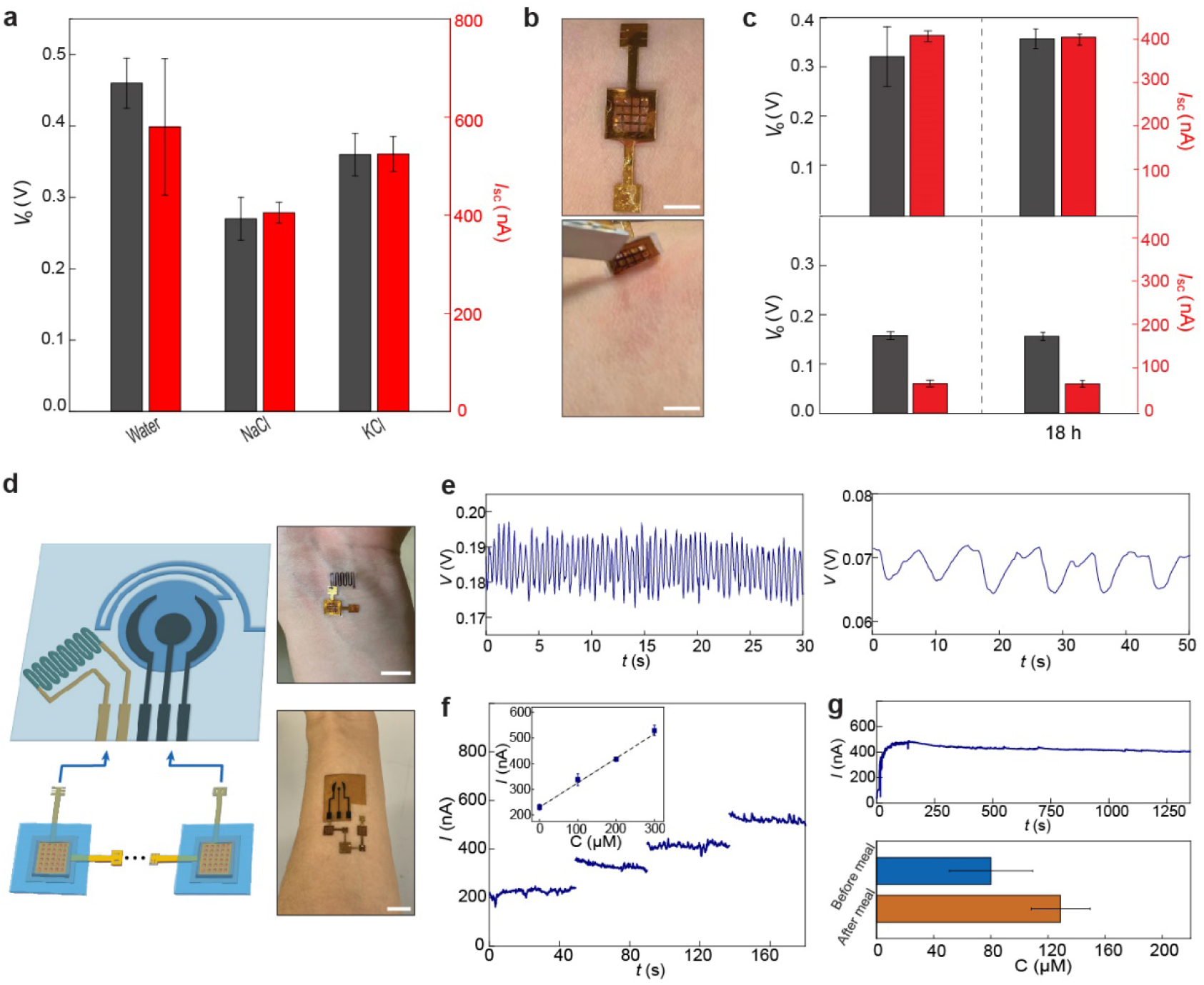
Wearable powering. **a**, Open-circuit voltage *V*_o_ (gray) and short-circuit current *I*_sc_(red) from biofilm devices placed on deionized water, 0.5 M NaCl, and 0.5 M KCl solutions. **b**, A biofilm device patched on skin (top) and being removed 18 h later (bottom). Scale bars, 0.5 cm. **c**, *V*_o_(gray) and *I*_sc_ (red) from biofilm devices patched on sweating skin (top) and dry skin (bottom), before (left) and after (right) 18 h. **d**, (Left) schematic of connecting biofilm devices to wearable sensors for wearable powering. (Right) Actual photos of powering a skin-wearable strain sensor with one biofilm device (top) and an electrochemical glucose sensor with three biofilm devices (bottom). Scale bars, 1 cm. **e**, Measured pulse signal (left) from the wrist and respiration signal (right) from the chest using the biofilm-powered strain sensor. **f**, Amperometric responses from a biofilm-powered glucose sensor placed in solutions having glucose concentration (*C*) of 0, 100, 200, and 300 µM, respectively. The inset shows the calibrated response curve. **g**, (Top) A continuous measurement of current from a biofilm-powered glucose sensor during exercise. (Bottom) Calibrated glucose levels from collected measurements before (blue) and after (orange) a meal.

The device with the PDMS seal (Fig. 2a) facilitated skin adhesion and the porous biofilm provided a breathable interface, so that prolonged (>12 h) contact did not irritate the skin. A device patch worn on sweaty skin produced power comparable to that produced with the salt solution and maintained its performance after 18 h (Fig. 3c, top). Even non-sweating skin generated a substantial electric output (Fig. 3c, bottom), demonstrating that a continuous low-level secretion of moisture from skin is sufficient to drive this hydroelectric output.^21-23^ Thus, the biofilm-sheet device is a promising candidate for continuous powering of wearable electronics.

As proof-of-concept skin-wearable biofilm-sheet devices were connected to wearable sensors (Fig, 3d). These included a strain sensor for self-powered monitoring of pulse (Fig. 3e, left), respiration (Fig 3e, right), and other bodily signals (Fig. S13). Interconnected biofilm-sheet devices were able to power a laser-patterned^24,25^ wearable electrochemical glucose sensor (Fig. 3f; Fig. S14) that monitored the glucose in sweat during exercise (Fig. 3g, top panel), and differentiated glucose levels before and after eating (Fig. 3g, bottom panel).

The previously recognized mechanism for electricity generation with biofilms of electroactive microbes like *G. sulfurreducens* is their ability to oxidize organic substrates with electron transfer to electrodes in microbial fuel cells.^4^ However, this technology requires that the microbes be maintained in a viable state under anoxic conditions with a continuous supply of organic fuel. In contrast, the biofilm-sheet devices achieved comparable or better energy densities and a faster start-up time^10^ while not dependent on cell viability. Storing the biofilms in air for more than a month or baking them at 90 °C had no impact on current generation (Fig. S4c; Fig. S15).

The reason that the biofilms-sheet devices generate more power than wood-based systems^12^ for evaporation-based electricity generation is probably related to the high surface area of hygroscopic and charged moieties within the biofilm,^16^ coupled with abundant channels for fluid flow, that are associated with the spacing of small microbial cells in the biofilm matrix (Fig. 1c). These features enhance both water and charge transport. The estimated surface area within the biofilm (∼8 m^2^/cm^3^) is much greater than that of wood or carbon foam.^14,26^ Films assembled from nanomaterials of smaller sizes,^12,27^ lack a supportive matrix and hence have tight physical contact between individual nanoelements.

An alternative approach to growing biofilms may be to collect planktonic cells as a mat on a filter. This approach was previously explored for making films of *G. sulfurreducens* protein nanowires for electricity production from atmospheric humidity.^28^ Devices for evaporation-based power generation fabricated with mats of filtered *G. sulfurreducens* (Fig. 4a; Figs. S16) generated electricity (Fig. 4b, middle and right) with an energy output ∼30 % that of the same size biofilm-sheet devices (Fig. 2b). To determine if evaporation-based electricity generation is possible with mats of other microbes, mats were fabricated with *Escherichia coli*, a microbe amenable to mass cultivation. The *E. coli* mats were less effective in energy output (∼15 %) than the *G. sulfurreducens* mats in current generation. Transmission electron microscopy revealed that, due to the lack of an extracellular polymeric matrix, the porosity of the filtered mats (Fig. 4d) was much lower than in the biofilm-sheets (Fig. 1d), which is expected to reduce fluid and charge transport.

**Figure 4.**
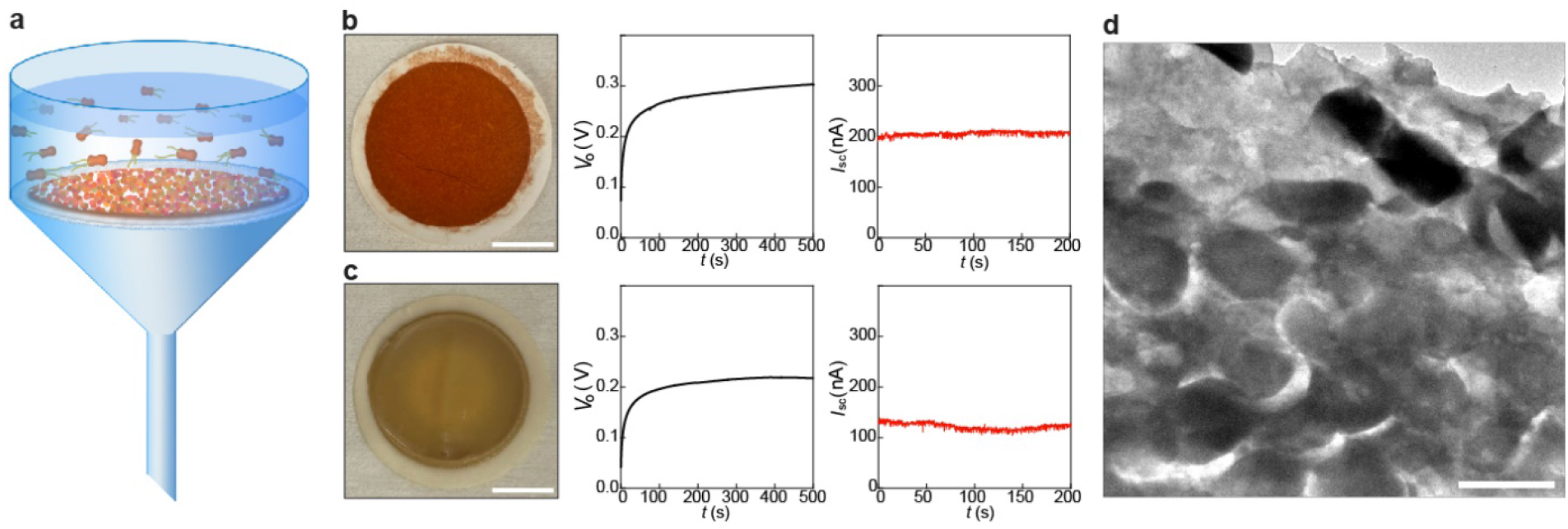
Devices made from filtered cells. **a**, Schematic of harvesting cells from microbial solution by filtering. **b**, Representative *V*_o_ (middle) and *I*_sc_ (right) measured from a device made from filtered *G. sulfurreducens* cells (left). Scale bar, 1 cm. **c**, Representative *V*_o_ (middle) and *I*_sc_(right) measured from a device made from filtered *E. coli* cells (left). Scale bar, 1 cm. Both devices in (b) and (c) had the same structure as shown in Fig. 2a with a size of 5×5 mm^2^. **d**, Cross-sectional TEM image of a *E*.*coil* mat assembled by filtering planktonic. Scale bar, 1 µm.

These results demonstrate that biofilm-sheets are a novel, sustainably produced material capable of scalable power production from evaporation-based electricity generation. Other strategies for organizing microbial cells into highly channelized, high surface area materials may be feasible. The ubiquity of microorganisms and their proclivity for biofilm formation suggests possibilities for harvesting electricity via similar evaporation-based strategies in diverse environments.

## Methods

### Biofilm preparation

The conductive biofilms of *G. sulfurreducens* strains CL-1 were routinely cultured as previously described.^15^ Briefly, the biofilms were grown anaerobically in two-chambered H-cell systems with a continuous flow of medium with acetate (10 mM) as the electron donor and a polished graphite stick anode poised at 300 mV versus Ag/AgCl as the electron acceptor. For studies with mats of cells collected with filters, wild-type *G. sulfurreducens* was grown as previously described^29^ under anaerobic conditions with acetate as the electron donor and fumarate as the electron acceptor. Planktonic cells of *E. coli* were grown in LB medium as previously described.^30^ Planktonic cells were filtered through a filter paper (42.5 mm dia., 8 µm pore size; Whatman) (Supplementary Fig. S16). The filtered mat of cells was then washed with deionized (DI) water.

### Fabrication of biofilm devices

The biofilms were patterned using a laser writer (Supplementary Fig. S3) and subsequently transferred onto substrates for device fabrication. Fabrication details for the planar single device (Fig. 1b-i) and array device (Fig. 1b-ii) can be found in Supplementary Fig. S4 and Fig. S8, respectively. The mesh electrode was fabricated by coating a thin gold layer on a laser-patterned polyimide (PI) mesh scaffold (Supplementary Fig. S9).

### Imaging

In order to examine cross sections of biofilms with transmission electron microscopy (TEM) biofilms were fixed (2% paraformaldehyde and 0.5% glutaraldehyde in 50 mM 1, 4-piperazinebis (ethanesulfonic acid) (PIPES) at pH 7.2) for 1 h at room temperature and washed 3 times with 50 mM PIPES. The biofilm samples were then dehydrated with a graded ethanol series (30, 50, 70, 80, 100%; 30 min each stage with gentle agitation). The dehydrated samples were infiltrated with LR White (medium grade, Electron Microscopy Sciences) and polymerized at 55 ºC overnight. Thin sections (∼50 nm thick) of fixed biofilm were made with a microtome (ULtracut S; Leica Microsystems). Sections were positively stained with 2% uranyl acetate and imaged (FEI Tecnai-T12 TEM) at 80 KV. The thickness of biofilms was acquired using 3D profiler (NewView™ 9000; Zygo).

### Fabrication of glucose sensors

The carbon electrodes were defined by a laser writer (LaserPro Spirit GLS; GCC) on a polyimide substrate.^24,25^ An Ag layer was electrodeposited onto the reference electrode by immersing it in a plating solution (250 mM silver nitrate, 750 mM sodium thiosulfate and 500 mM sodium bisulfite), applied with −0.2 mA for 100 s. The rest electrodes were electrochemically deposited with a Pt layer, by immersing them in a solution (1.0 mM H_2_PtCl_6_ dissolved in 0.5 M H_2_SO_4_) and scanning the electrodes from + 0.5 to − 0.7 V for consecutive 30 cycles at 100 mV/s. The glucose oxidase (Gox) enzyme was subsequently immobilized by dip-coating the Pt electrode in a solution containing 140 mg/ml Gox, 56 mg/ml bovine serum albumin, and 25 w% glutaraldehyde, followed by a 2-h soak in phosphate buffered saline to remove residue. A double-sided medical adhesive (MH-90445Q; Adhesives Research), defined with microfluidic channel (200 µm wide) and reservoir (5 mm diameter) by the laser writer, was attached to the PI substrate with defined electrodes. The sensing signals were acquired using an electrochemical workstation (CHI 440; CH Instruments).

### Fabrication of strain sensors

The serpentine electrode (0.5 mm wide, 1 cm long, Cr/Pt = 10/20 nm) was defined on a polyimide film by standard photolithography, metal evaporation, and lift-off processes. Cracks were generated in the electrode by stretching the substrate (2%) using a mechanical testing system (ESM303; Mark-10 Inc.). The sensor was attached on the skin with a double-sided medical adhesive.

### Electrical measurements

Electrical measurements were performed in the ambient environment, unless otherwise specified. The biofilm devices in Fig. 1b-i were tested by immersing one end of the biofilm into a petri dish filled with DI water. The biofilm devices with mesh electrodes (Fig. 2a) were tested by placing them on wet tissue papers soaked with DI water. The voltage and current outputs were measured by using a Keithley 2401.

## Notes

The authors declare no competing financial interest.

## Acknowledgements

J.Y. and D.R.L. acknowledge support from the National Science Foundation (NSF) DMR2027102. X.L. acknowledges support from the Link Foundation Energy Fellowship. J.Y. also acknowledges supports from NSF CAREER CBET-1844904 and NSF ECCS-1917630. Part of the device fabrication work was conducted in the clean room of the Center for Hierarchical Manufacturing (CHM), an NSF Nanoscale Science and Engineering Center (NSEC) located at the University of Massachusetts Amherst.

## Author contributions

J.Y. and X.L. conceived the project and designed experiments. D.R.L. oversaw material design and production. X.L. carried out experimental studies in material characterization, device fabrication and electrical measurement. H.Y., T.F., S.F. and L.S. helped with device fabrication and characterization. T.U., T.L.W. and K.P.N. prepared biofilms and bacteria solution. J.Y., D.R.L. and X.L. wrote the paper. All authors discussed the results and implications and commented on the manuscript.

## Notes

### Competing Interest Statement

The authors have declared no competing interest.

