## Supplementary Information for "Generating Electricity from Water Evaporation Through Microbial Biofilms"

---

1. Department of Electrical Computer and Engineering, University of Massachusetts, Amherst, MA, USA.
2. Department of Microbiology, University of Massachusetts, Amherst, MA, USA.
3. Institute for Applied Life Sciences (IALS), University of Massachusetts, Amherst, MA, USA.
4. Department of Biomedical Engineering, University of Massachusetts, Amherst, MA, USA.

**This file includes:**

Supplementary Figures S1 – S16

Supplementary References

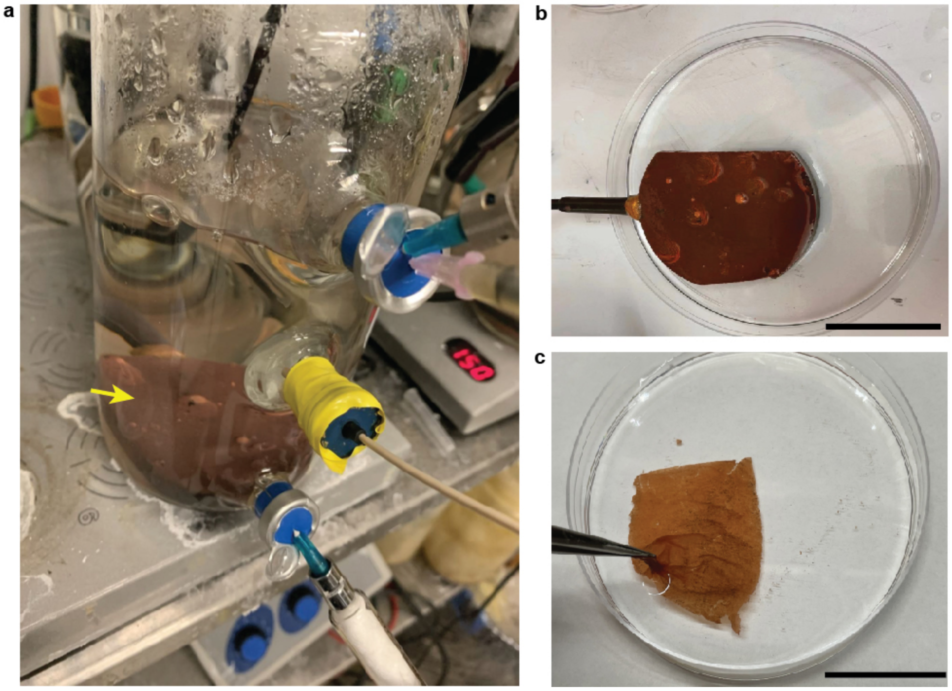

**Fig S1**. **Biofilm growth and harvesting**. **a**, Setup for *G. sulfurreducens* biofilm growth. The arrow indicates the growth electrode. Details about the growth process can be found in *Methods*.[1] **b**, As-grown *G. sulfurreducens* biofilm on a polished graphite electrode. Scale bar, 5 cm. **c**, A harvested biofilm floating on water. The biofilm was harvested by scraping it off the electrode with a blade, rinsed in water, and stored in a 4 °C refrigerator. Scale bar, 5 cm.

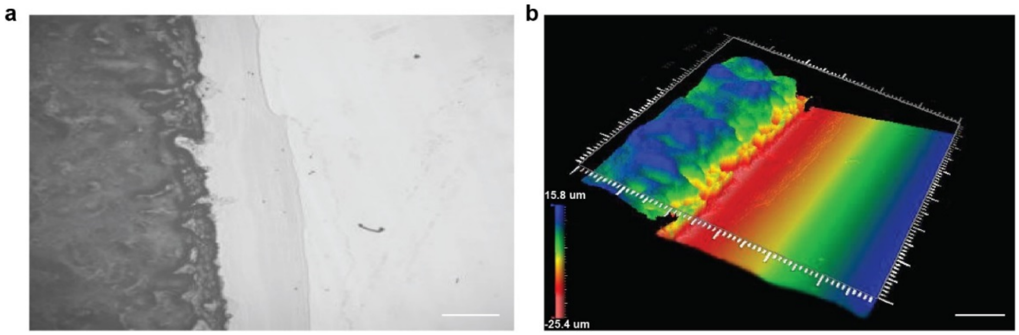


**Fig. S2**. **Measurement of biofilm thickness**. **a**, Optical image of a *G. sulfurreducens* biofilm (dark region) prepared on a silicon substrate. Scale bar, 1 mm. **b**, 3D topological image of the biofilm by using a non-contact 3D profiler (NewView™ 9000; Zygo). Scale bar, 1 mm. The measured thickness was 38 ± 3 µm. The biofilm was kept wet during the imaging to reflect the thickness under device working condition.

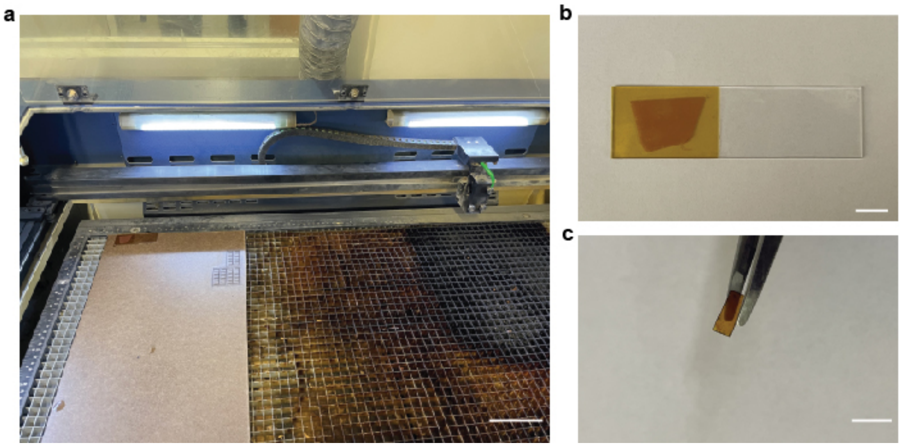


**Fig. S3**. **Biofilm patterning by laser writing**. **a**, The laser writing system (LaserPro Spirit GLS; GCC). Scale bar, 10 cm. **b**, An as-prepared *G. sulfurreducens* biofilm before patterning. The biofilm was placed on polyimide (PI) substrate further supported by a glass slide (for easy transfer). Scale bar, 1 cm. **c**, A patterned biofilm by the laser writer (15% power, Speed 20%, 400 points per inch resolution). Scale bar, 1 cm.

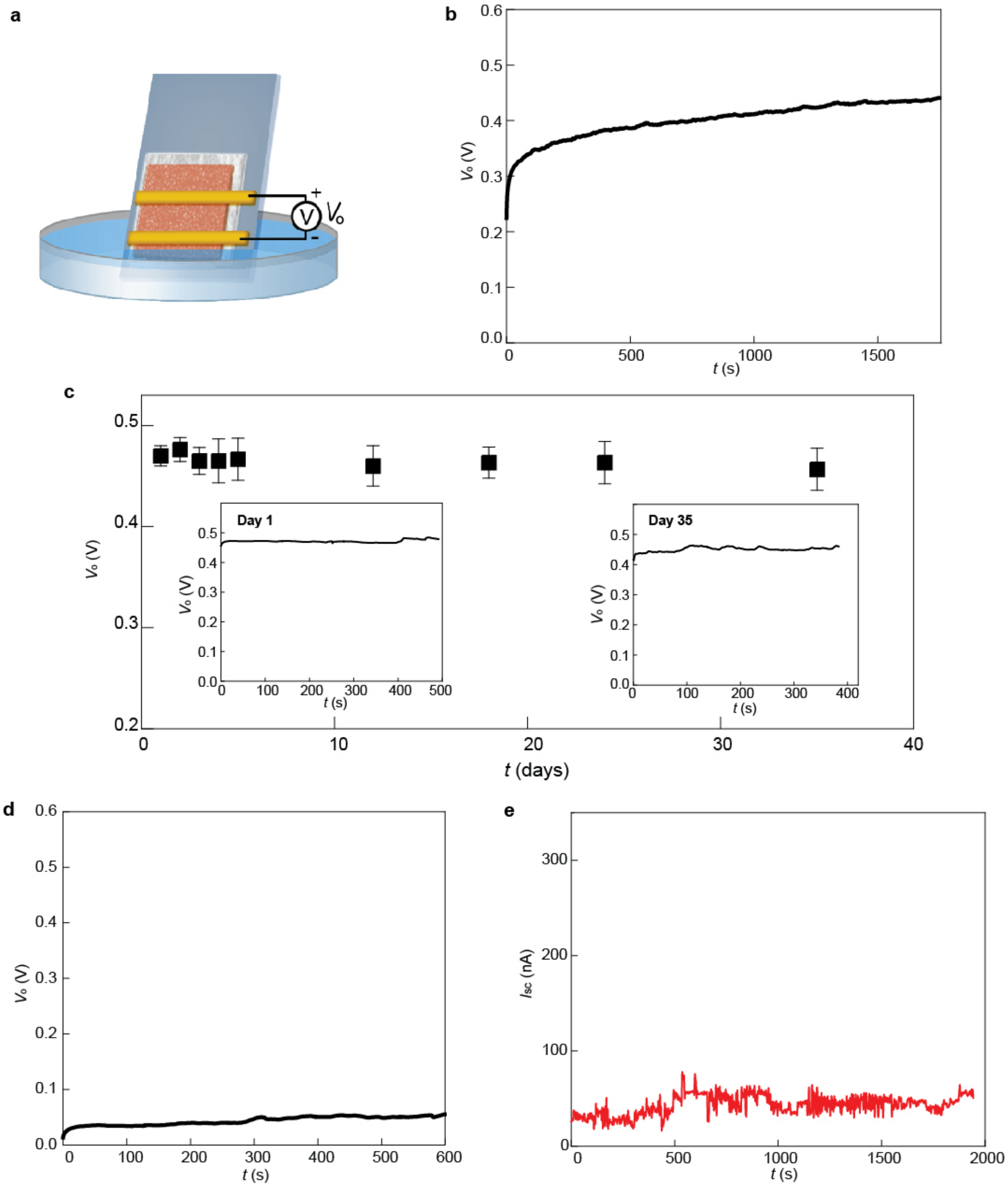


**Fig. S4**. **Electric output from *G. sulfurreducens* biofilm device and control**. **a**, Schematic of the biofilm device used in Fig. 1d in the main text. The biofilm was placed on a tissue paper supported by a glass slide (25 × 75 mm$^2$). A pair of electrodes (2 × 15 mm$^2$), made from Au-coated polyimide films, was covered on top of the biofilm. The tissue paper facilitated the water transport but did not contribute substantially to energy output as shown in (d) and (e). Devices without tissue paper produced the same effect (Fig. S10). **b**, The measured open-circuit voltage ($V_o$) from a biofilm device. The size of the biofilm device was 2 × 10 mm$^2$. **c**, A continuous 35-day recording of the $V_o$ from the device. The insets show representative recording curves on day-1 and day-35, respectively. **d**, Voltage output ($V_o$ ~0.05 V) from the tissue paper after removing biofilm. **e**, Short-circuit current output ($I_{sc}$ ~30 nA) from the tissue paper after removing the biofilm. The measurements were performed in the ambient environment with a relative humidity (RH) of ~50%.

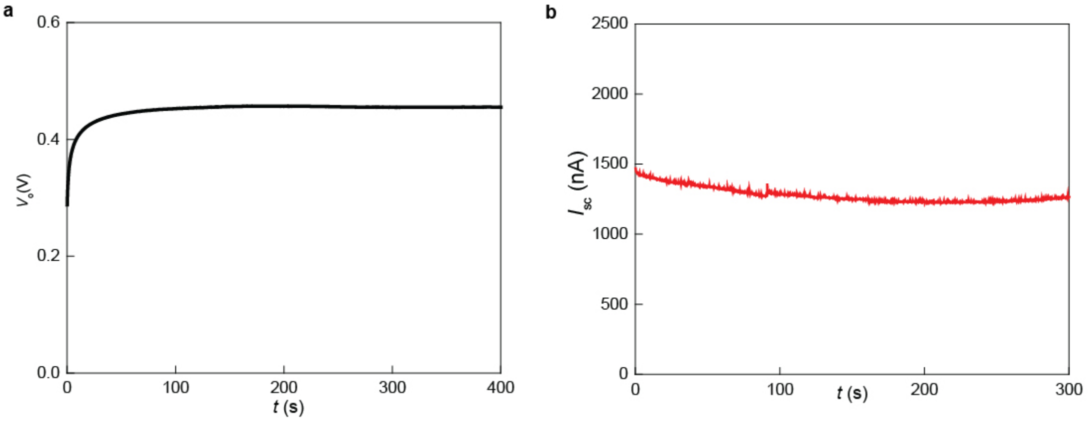


**Fig. S5**. **Biofilm device using carbon electrodes**. **a**, Open-circuit voltage ($V_o$) measured from a *G. sulfurreducens* biofilm device having the same structure as shown in Fig. S4, with the pair of Au electrodes replaced by a pair of carbon electrodes. The carbon electrodes were defined by printing conductive carbon ink on polyimide stripes (Dimatix Inkjet DMP 2831; Fujifilm). **b**, Measured short circuit current ($I_{sc}$) of the device. The measurements were performed in the ambient environment with a RH ~50%. Both results were close to values obtained with Au electrodes.

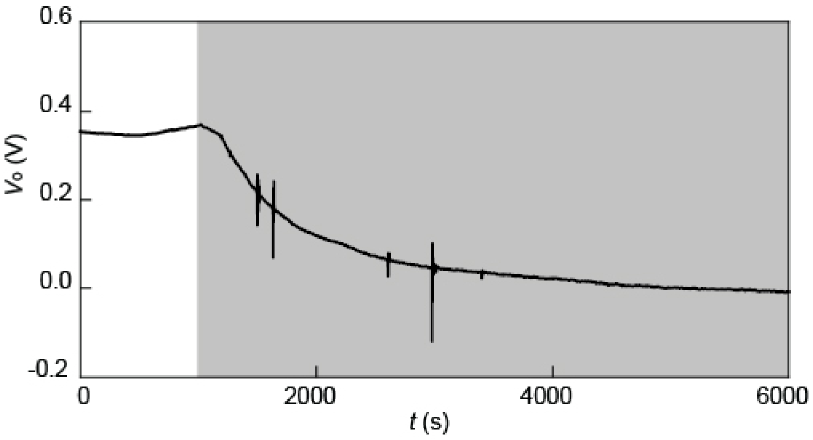


**Fig. S6**. Voltage output ($V_o$) from a biofilm device during the depletion of water source (gray region). The device had the same structure as shown in Fig. S4.

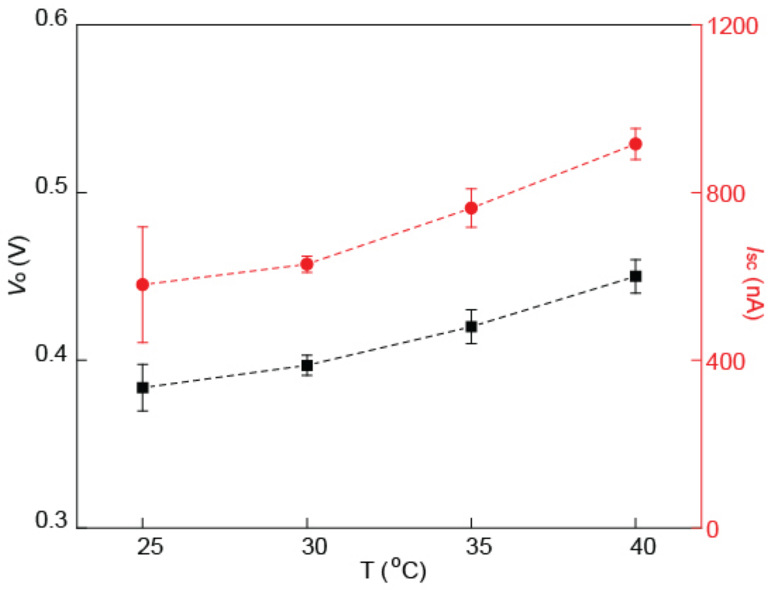


**Fig. S7**. **Measured open-circuit voltage ($V_o$) and short-circuit current ($I_{sc}$) from a biofilm device at increasing water evaporation**. The temperature was increased from 25 to 40 ℃ to increase the water evaporation rate. To improve the uniformity in temperature across the device/water interface, a mesh-electrode biofilm device (Fig. 2a in main text; size ~5×5 mm$^2$, porosity 0.4) was used. The device was placed on a tissue paper soaked with water, which was then placed on a hot plate held at the specified temperatures.

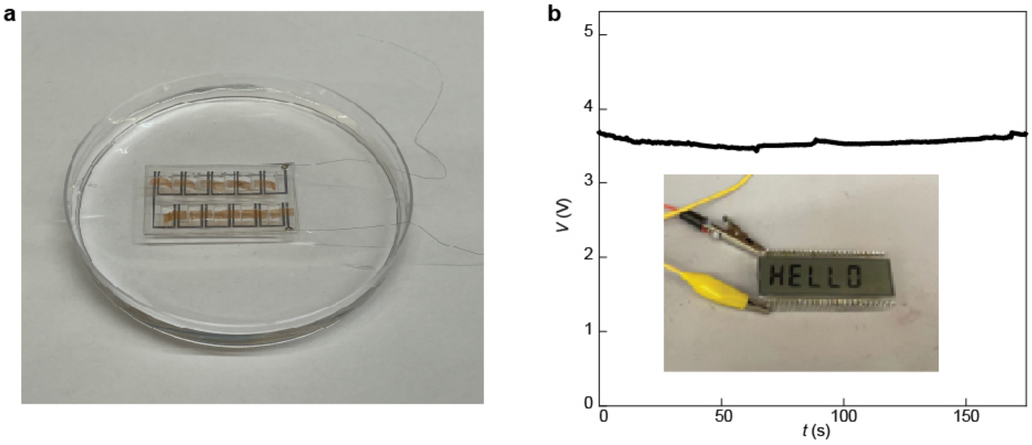


**Fig. S8**. **Interconnected biofilm device array**. **a**, 10 biofilm devices were connected in series on a polyethylene terephthalate (PET, ~80 μm thick) substrate, with the PET substrate further placed on a polydimethylsiloxane (PDMS, ~0.5 mm thick) substrate floating on water (device structural schematic can be found in Fig. 1b-ii in the main text). The biofilms and openings on PET substrate were patterned with a laser writer (LaserPro Spirit GLS; GCC). The spacing of each pair of electrodes was 1 mm and the size of each biofilm was ~3×7 mm$^2$. **b**, Voltage output (*V*) measured from the device array, which was used to power a LCD (VI-602-DP-FC-S; Varitronix) as shown in inset.

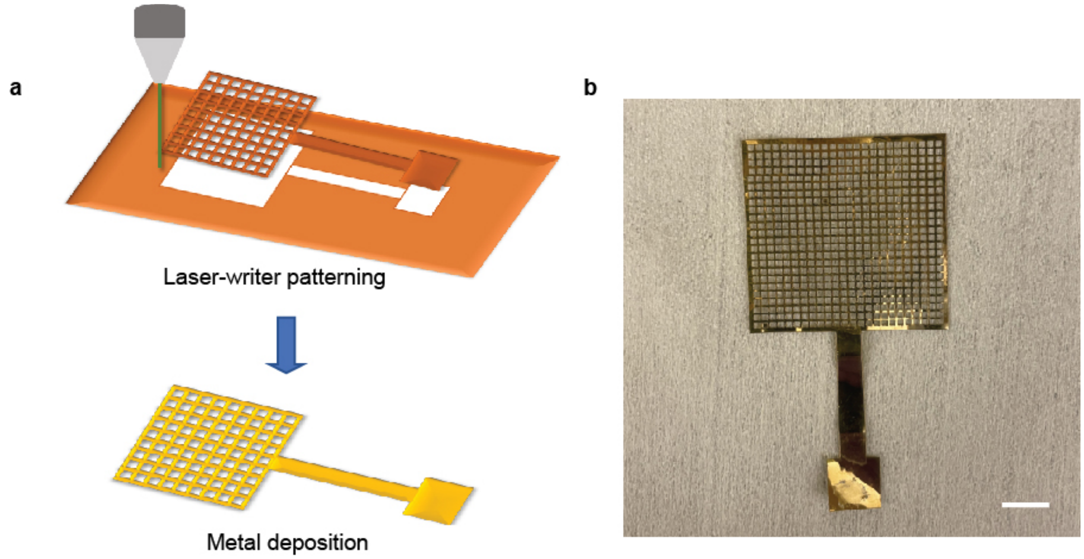


**Fig. S9**. **Fabricating mesh electrode**. **a**, Schematics of fabricating the mesh electrode, involving (top) patterning a PI substrate using a laser writer (LaserPro Spirit GLS; GCC) and (bottom) coating the defined PI mesh with a metal layer (Cr/Au = 5/50 nm) using standard metal deposition. **b**, An as-fabricated mesh electrode. Scale bar, 5 mm.

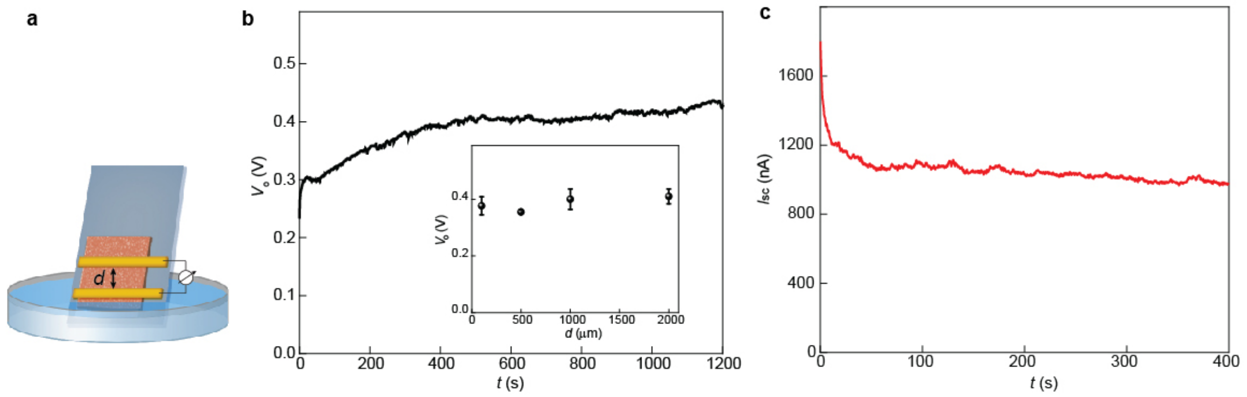


**Fig. 10**. **Electric outputs from biofilm devices of different electrode spacings**. **a**, Schematic of the biofilm device. A biofilm (5×10 mm$^2$, without tissue paper support) was first placed on a glass substrate (25×75 mm$^2$), with its one end immersed in water. A series of pairs of electrodes with spacings (*d*) of 100, 500, 1000, and 2000 μm were placed on the biofilm to measure the electric outputs. The electrodes were made from Au-coated PI films (2×15 mm$^2$). **b**, A representative open-circuit voltage ($V_o$) measured with a pair of electrodes spaced ~100 μm. (Inset) Measured $V_o$ at different electrode spacings. The trend shows that $V_o$ saturated at sub-100 μm spacing, which is consistent with the observation that biofilms of sub-100 μm thickness generated similar voltage across the vertical thickness (Fig. 2 in main text). **c**, Short-circuit current ($I_{sc}$) measured from the device with the same pair of electrodes spaced ~100 μm. The measurements were performed in the ambient environment with a RH of ~50%.

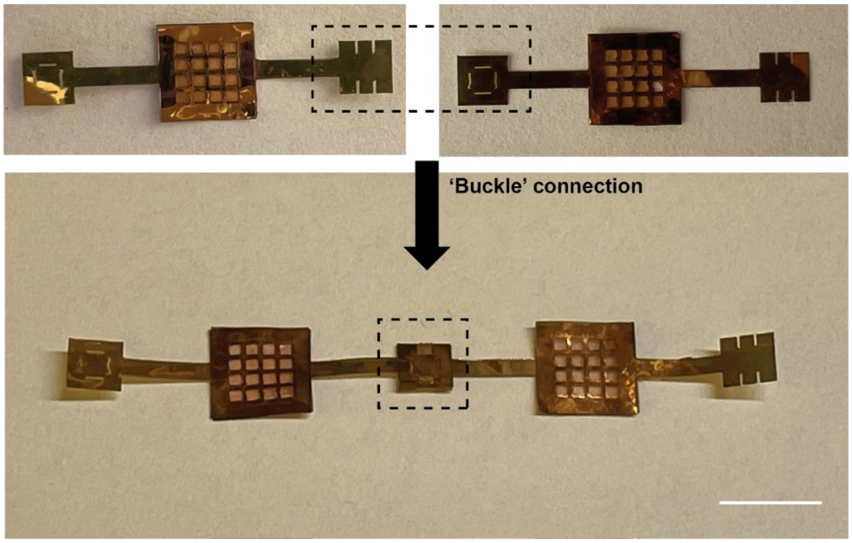


**Fig. S11**. **The design of a 'buckle' connection**. The contacts of top and bottom *mesh* electrodes were patterned with socket and plug features by laser writing as shown in the top panel, which enabled an easy buckle connection between the devices (bottom). Note that the design also allowed for a 90-degree rotation in the connection. Scale bar, 5 mm.

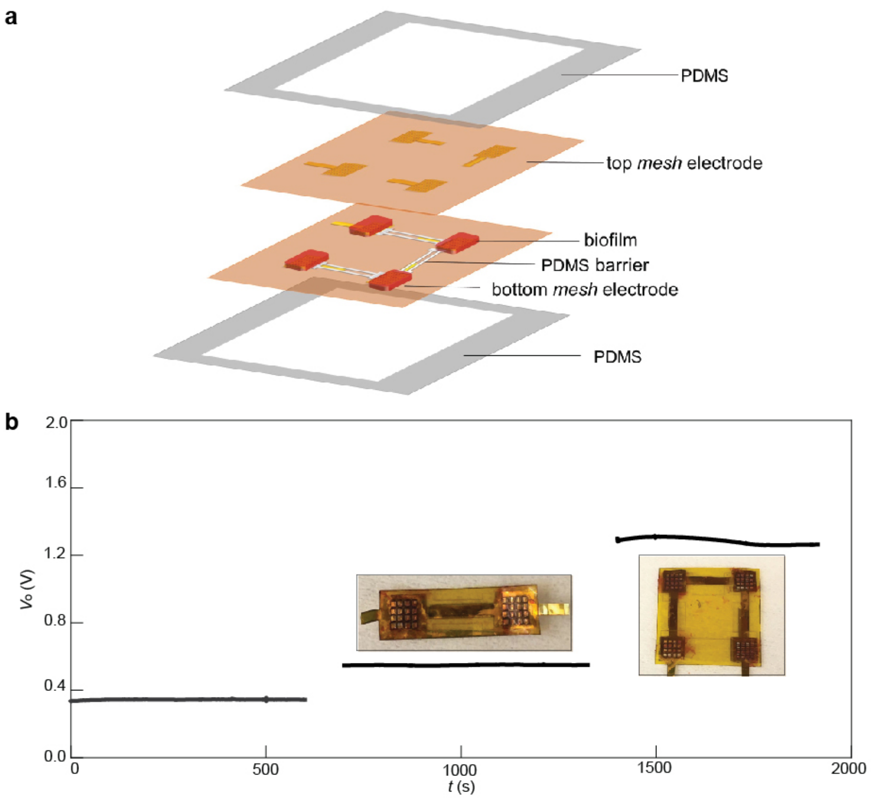


**Fig. S12**. **On-substrate integration of biofilm devices**. **a**, Schematic of series integration of biofilm devices. The top and bottom *mesh* electrodes were defined on polyimide films using a laser writer, followed by selected-area metal deposition (Cr/Au = 5/50 nm) using a shadow mask. The contacts of top and bottom mesh electrodes were arranged in a way that they would form series connection when put into contact. Patterned biofilms were then transferred onto the bottom mesh electrodes. A molded PDMS barrier layer (~20 µm thick) was sandwiched between the top and bottom electrodes to prevent short circuit at the biofilm edge. The device was then sealed by two PDMS layers (~20 ×µm thick). **b**, Open-circuit voltage ($V_o$) measured from fabricated devices containing 1, 2, and 4 biofilms, showing linear increase in output.

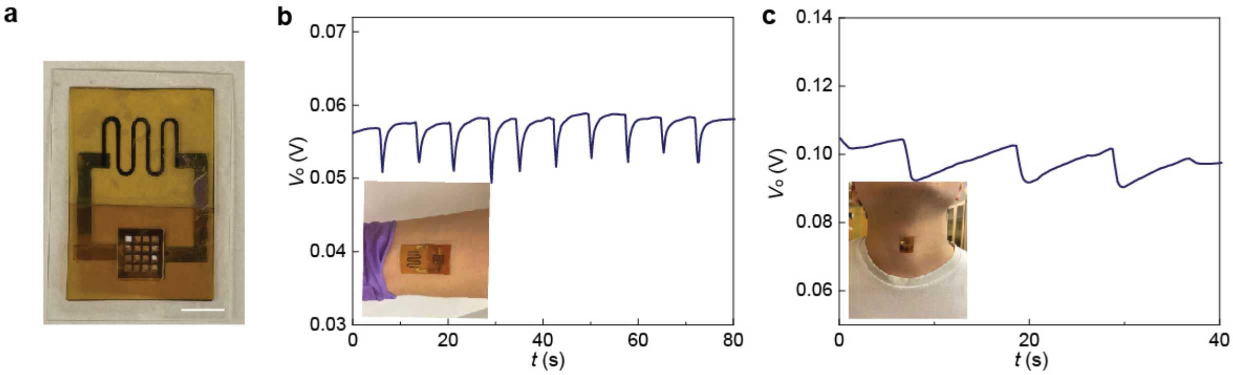


**Fig. S13**. **Integrated sensor and biofilm for wearable detection**. **a**, An integrated wearable patch with a crack strain sensor[2] directly connected to the biofilm device. Details of fabrication can be found in *Methods*. Scale bar, 5 mm. **b**, Measured bodily mechanical signal when bending the wrist using the wearable patch. **c**, Measured bodily mechanical signal of swallowing using the same patch. The signals were acquired by connecting the two terminals of the crack sensor to a source meter (Keithley 2401; Keithley Instruments).

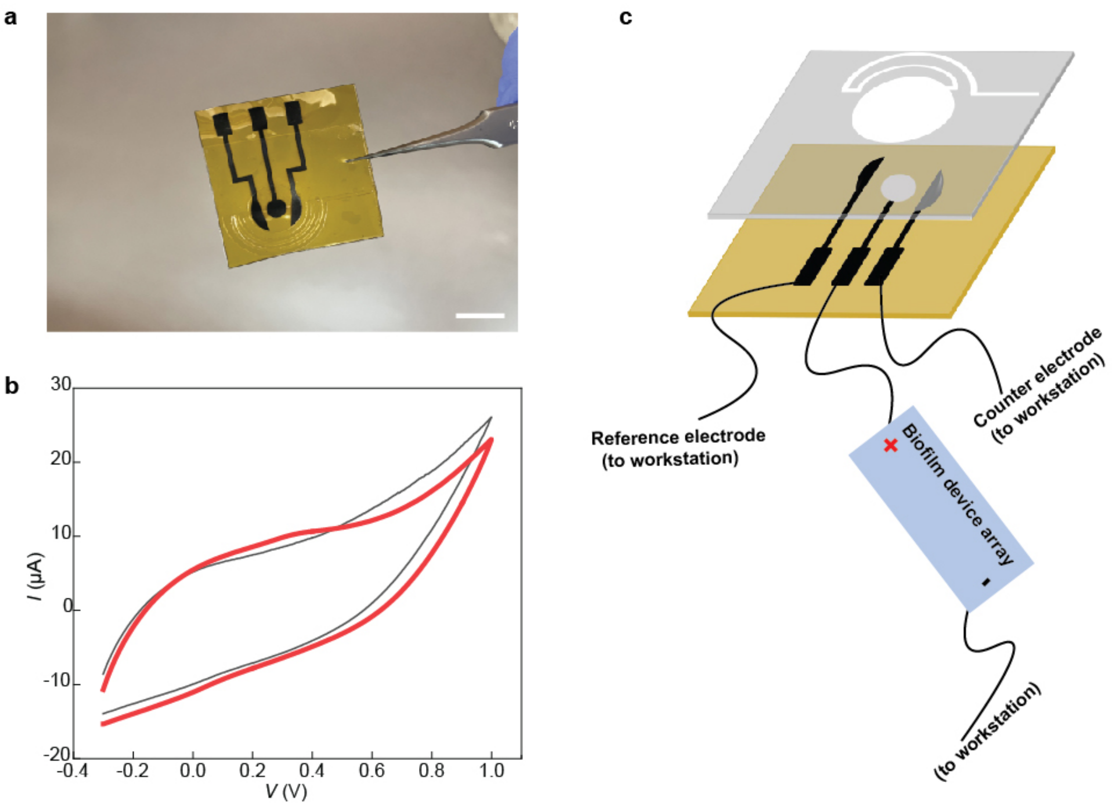


**Fig. S14**. **Biofilm-powered glucose sensor**. **a**, The glucose sensor consisted of laser-induced carbon electrodes on a polyimide substrate,[3,4] covered with a medical tape (MH-90445Q, Adhesives Research, Inc) with an engraved opening. Scale bar, 5 mm. Details of fabrication and functionalization processes can be found in *Methods*. **b**, Cyclic voltammetry (CV) measurement was first performed to identify the oxidization threshold voltage. CV from 500 μM glucose solution (red) showed oxidization peak ~0.4 V compared to reference (gray) from phosphate buffered saline solution without glucose. **c**, For real-time sensing, biofilm devices were connected to the working electrode to provide an effective voltage larger than 0.4 V. The three terminals were connected to an electrochemical workstation (CHI 440; CH Instruments) to acquire the current signal.

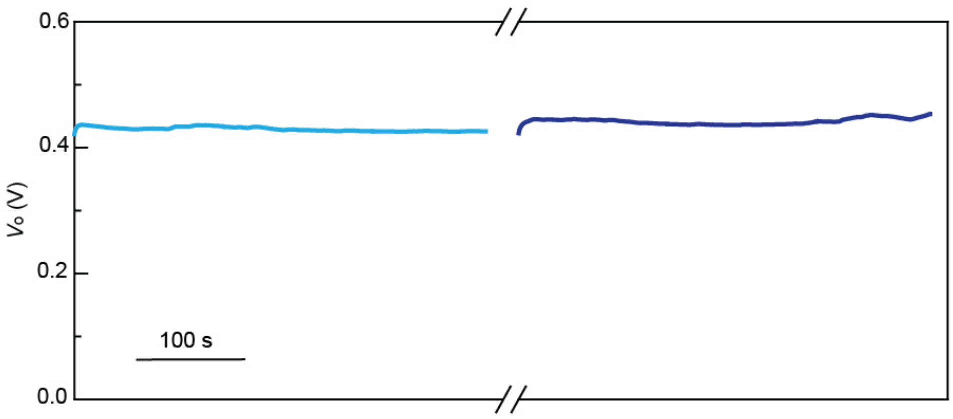


**Fig. S15**. Open-circuit voltage ($V_o$) measured from a biofilm device before (light blue) and after (dark blue) heating to 90°C for 30 min. The device had the same structure as in Fig. S4.

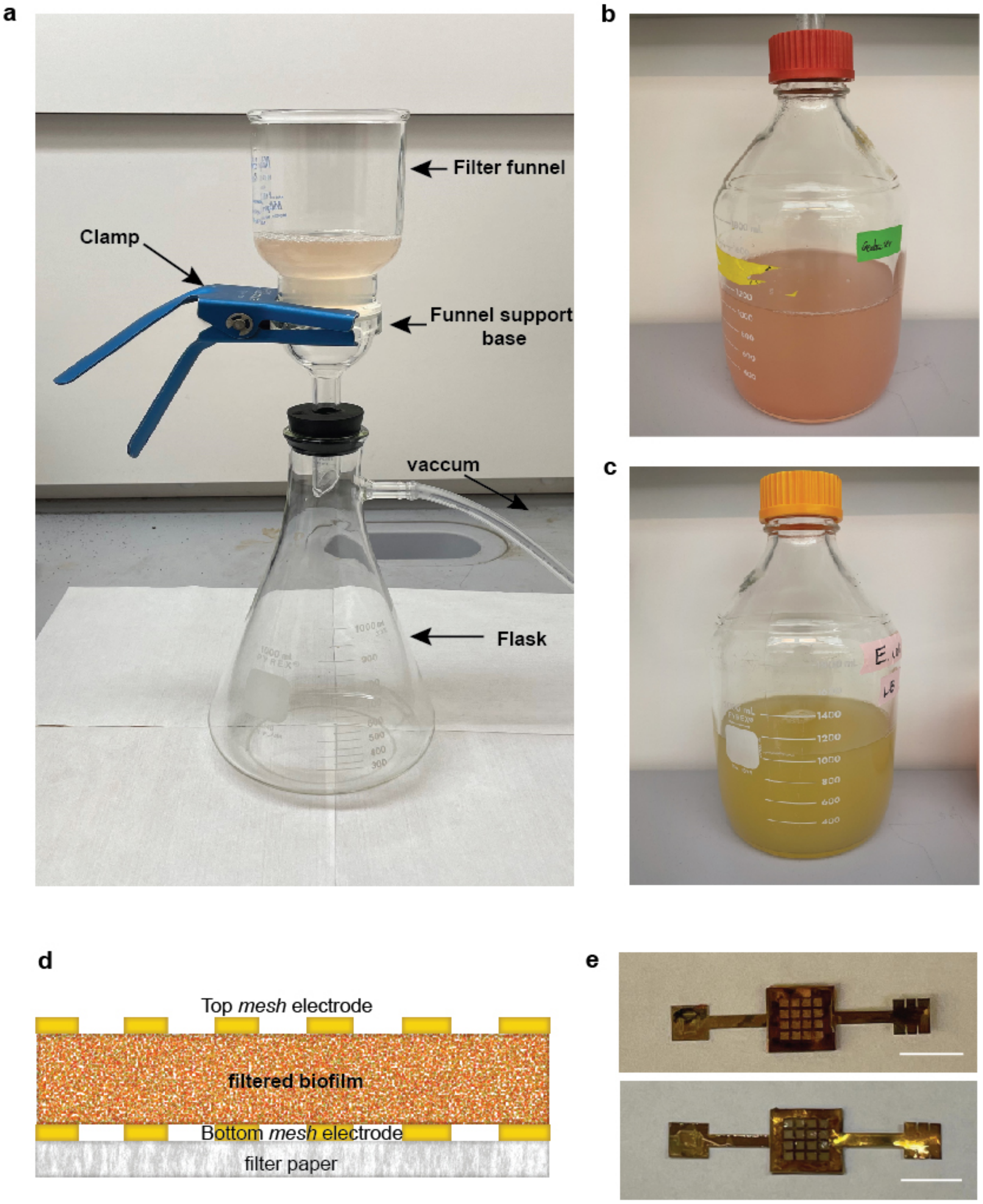


**Fig. S16**. **Mats of filtered cells and devices**. **a**, The setup of the filtering system. The microbial solution was passed through a papers filter (42.5 mm dia., 8 µm pore; Whatman) with vacuum filtration. A 300 ml volume was filtered repeatedly until the solution appeared to be clear. **b**, Prepared *G. sulfurreducens* solution for filtration. **c**, Prepared *Escherichia coli* solution for filtration. Details of the growth and solution preparation can be found in *Methods*. **c**, Cross-sectional schematic of the device made from filtered cells. The bottom mesh electrode was first placed on the filter paper before filtration. After filtration the top mesh electrode was capped. The filter paper was cut into the same size of the mesh electrode and left in the device, which would be in direct contact with water and hence did not block water evaporation. **e**, Fabricated devices using filtered biofilms of (top) *G. sulfurreducens* and (bottom) *E. coli*. Scale bars, 5 mm.
